# Dendritic spikes expand the range of well-tolerated population noise structures

**DOI:** 10.1101/454215

**Authors:** Alon Poleg-Polsky

## Abstract

The brain operates surprisingly well despite the noisy nature of individual neurons. The central mechanism for noise mitigation in the nervous system is thought to involve averaging over multiple noise-corrupted inputs. Subsequently, there has been considerable interest recently to identify noise structures that can be integrated linearly in a way that preserves reliable signal encoding. By analyzing realistic synaptic integration in biophysically accurate neuronal models, I report a complementary de-noising approach that is mediated by focal dendritic spikes. Dendritic spikes might seem to be unlikely candidates for noise reduction due to their miniscule integration compartments and poor averaging abilities. Nonetheless, the extra thresholding step introduced by dendritic spike generation increases neuronal performance for a broad category of computational tasks, including analog and binary discrimination, as well as for a range of correlated and uncorrelated noise structures, some of which cannot be adequately resolved with averaging. This property of active dendrites compensates for compartment size constraints and expands the repertoire of brain states and presynaptic population activity dynamics can be reliably de-noised by biologically-realistic neurons.

**Significance Statement:** Noise, or random variability, is a prominent feature of the neuronal code and poses a fundamental challenge for information processing. To reconcile the surprisingly accurate output of the brain with the inherent noisiness of biological systems, previous work examined signal integration in idealized neurons. The notion that emerged from this body of work is that accurate signal representation relies largely on input averaging in neuronal dendrites. In contrast to the prevailing view, I show that de-noising in simulated neurons with realistic morphology and biophysical properties follows a different strategy: dendritic spikes act as classifiers that assist in extracting information from a variety of noise structures that have been considered before to be particularly disruptive for reliable brain function.

## Introduction

Neurons generate variable responses to the repetitive presentation of the same stimulus (Faisal et al., 2008). This variability, or noise, can corrupt the neuronal representation of the signal and limit the amount of usable information in neuronal populations (Kohn et al., 2016). Numerous models have examined the function of neural networks in noisy conditions [e.g. (Faisal et al., 2008; Franke et al., 2016; Kohn et al., 2016; Zylberberg et al., 2016)]. In virtually all studies, neurons were formulated to integrate their synaptic inputs linearly (McCulloch and Pitts, 1990). Linear integration is more likely for inputs that are distributed and activated in a sparse manner (Jia et al., 2010). In this regime, synaptic inputs are largely independent of each other and summate at the cell body.

However, most biological neurons are not linear integrators of synaptic inputs. Instead, dendritic voltage-dependent channels can dramatically affect synaptic processing. For example, numerous studies have demonstrated that dendrites can generate spikes in response to synaptic stimulation (Spencer and Kandel, 1961; Schiller et al., 2000; London and Hausser, 2005; Branco and Hausser, 2010; Major et al., 2013). The basic nonlinearity that results from dendritic spikes can underlie dynamic dendritic computational compartments, in which each dendritic unit integrates its inputs linearly and passes them through a nonlinear all-or-none filter (Mel, 1993; Poirazi et al., 2003; Polsky et al., 2004; London and Hausser, 2005; Major et al., 2013). Sufficiently strong activation of spatiotemporally coordinated synaptic inputs produces a regenerative dendritic spike that amplifies the synaptic drive to the dendrite and potentially boosts representation of salient input patterns (Lavzin et al., 2012; Smith et al., 2013; Palmer et al., 2014; Hawkins and Ahmad, 2016; Schmidt-Hieber et al., 2017).

Importantly, the size of the dendritic integration compartment for a dendritic spike can be surprisingly small; in many cell types, activation of only a handful of synaptic inputs is enough for generation of a dendritic spike (Major et al., 2008; Polsky et al., 2009; Branco and Hausser, 2011; Schmidt-Hieber et al., 2017). Although the concept of dendritic computational units is attractive, their small size may challenge their ability to accomplish reliable computations in the presence of a noisy synaptic drive.

Classically, the central neuronal denoising element has been assumed to be averaging over a large number of synaptic inputs (**Figure 1 B**). A rich theory emerged that divided noise structures between amenability and resistance to denoising by linear integration. However, a limitation of signal processing with dendritic spikes is that their integration compartments do not offer significant protection against high-variance noise. For this reason, trial-to-trial fluctuations that can be effectively negated by integration over multiple synaptic inputs in linear dendrites, are likely to have a significant impact on the reliability of computations in active dendrites.

**Figure 1.**
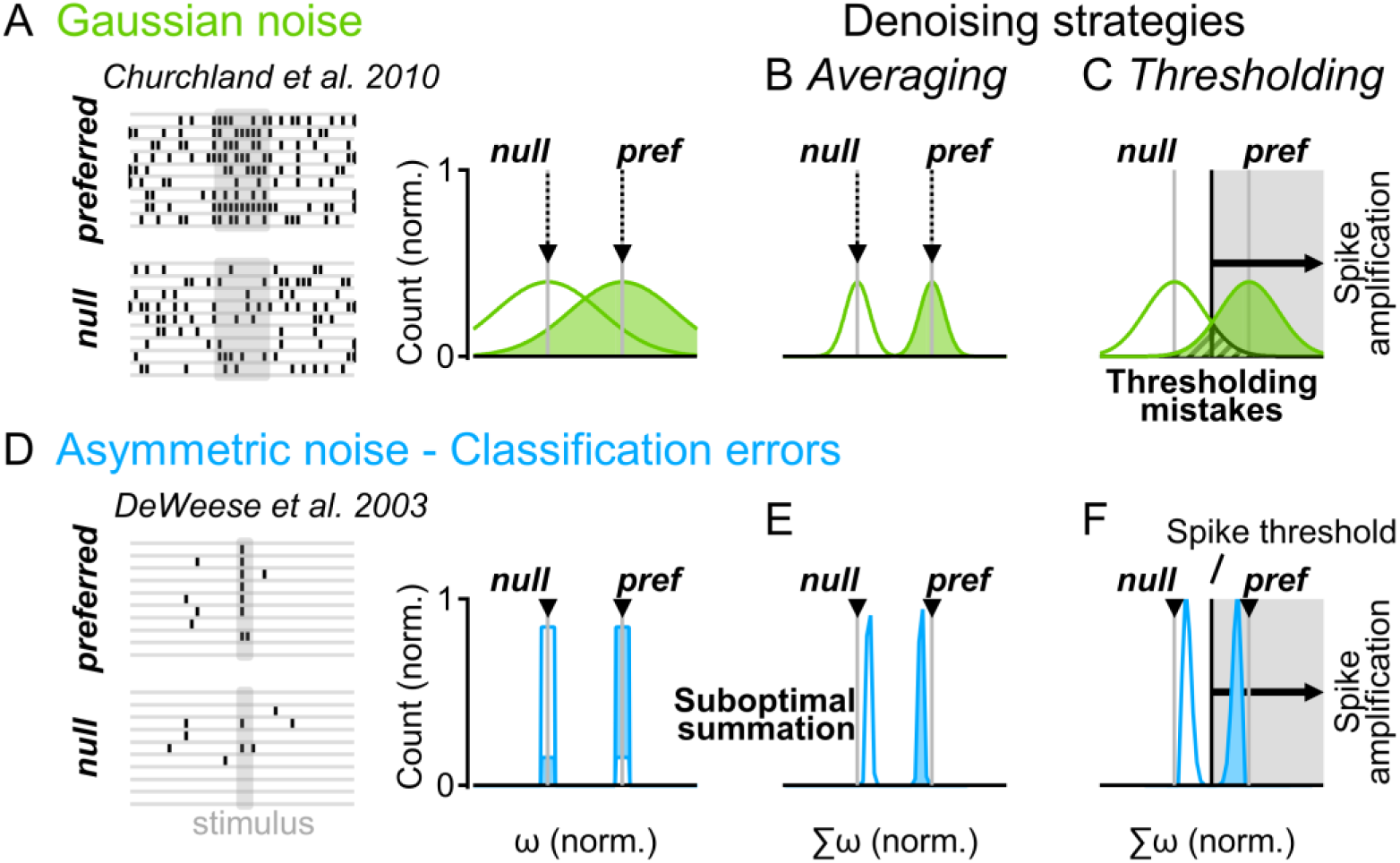
Distinct denoising operations in linear and active dendrites. (A) Representative spike trains (left) in a simulated neuronal population that respond to a stimulus with normally distributed firing rates (right). The mean firing rates (gray) correspond to noise-free responses. (B) Linear integration of a normally distributed population activity decreases the trial-to-trial variability of the postsynaptic cell. (C) The effectiveness of denoising by synaptic averaging depends on the size of the integration compartment. In active dendrites, dendritic spikes generation is typically governed by a small number of synaptic inputs, which limits the ability of the dendritic spike to reduce the negative impact of noise (‘thresholding mistakes’, black). (D-F) as in (A-C) for a simple asymmetric noise distribution of population responses, where preferred stimulation is signaled with a single spike (DeWeese et al., 2003). Some presynaptic cells produce ‘classification errors’ if they spike to the null or do not produce a response to the preferred stimulations (D). (E) Linear synaptic integration of an asymmetric presynaptic population shifts the mean output away from the noise-free population activity – diminishing the difference between stimuli representations (‘suboptimal summation’). (F) Spikes form a decision threshold that can help increase the separation between postsynaptic stimuli representations, compensating for the imperfect linear integration of asymmetric noises.

Here, I propose a distinct, and complementary role for dendritic subunits in information processing in the presence of noise. My theory is based on the observation that nonlinear dendritic spike generation enhances neuronal abilities to filter out noise based on a decision cutoff (**Figure 1 C**). To critically evaluate this concept, I first provide the conceptual rationale for how these denoising approaches apply to distinct input dynamics. Next, to demonstrate this effect in biophysically-realistic conditions, I test the prediction that active dendrites have unique processing capabilities in multi-compartmental models. My results demonstrate that dendritic units have an exceptional capability to mitigate noise structures that are paradoxically resistant to averaging, such as stimuli misclassification. Thus, active dendrites can compute in the presence of noise, and even surprisingly surpass the denoising capabilities of linear dendrites for a wide range of noise statistics. These outcomes hold for independent inputs, as well as for correlated presynaptic population dynamics. Taken together, the processing abilities of dendritic spikes challenge the noise tolerance rules formulated for linearly integrating neurons and extend the dynamic range of biological brains.

## Methods

Computer simulations were conducted on multicompartmental models using the NEURON 7.4 simulation platform. The modeled cell was subdivided into 483 compartments, with a maximum length of 19 µm. The area of the soma was 1350 µm^2^, and the total dendritic length was 3320 µm^2^. The resting membrane potential was set to -60 mV, the membrane resistance was set to 10,000 Ω·cm^2^, the axial resistance was 100 Ω·cm, and the membrane capacitance was set to 1 µF/µm^2^. Fast sodium channels (reversal potential=50 mV, gNa=200 mS/cm^2^), and delayed rectifier and slow non-inactivating potassium channels (reversal potential=-87 mV, gKdr=30 mS/cm^2^, gKs=1 mS/cm^2^) were used to allow for spike generation and adaptation, respectively (Lavzin et al., 2012).

### Synapses

Excitatory postsynaptic synapses contained AMPA (α-amino-3-hydroxy-5-methyl-4-isoxazolepropionic acid) Receptors (AMPARs) and NMDA (N-methyl-D-aspartic acid) Receptors (NMDARs). AMPAR currents had an instantaneous rise time and a decay time of 1.5 ms. NMDAR currents had a rise time of 2 ms and a decay time of 100 ms. The synaptic current reversed at zero, and the unitary AMPAR/NMDAR conductance was 1 nS. The NMDAR conductance voltage dependence was modeled as follows: 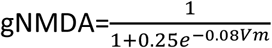 where Vm is the local membrane potential. When a linear synaptic integration was modeled, synapses were distributed over the entire dendritic tree and NMDAR conductance was typically set to zero to prevent spurious dendritic spike generation by randomly clustered synapses.

### Presynaptic stimulation

Directional simulations: The angle of preferred direction for each cell in the presynaptic population was chosen from a normal distribution with a mean at 180° and standard deviation (SD, σ) of 45°. The postsynaptic cell was assumed to prefer inputs from cells aligned with its preferred direction (PD)=180°; towards this goal, the amplitude of each presynaptic cell was multiplied by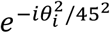, where *ϑi* is the difference (in degrees) between the PD angle of the presynaptic and the postsynaptic cell *i*. DSI was calculated as 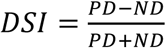 The DSI of the presynaptic population was set to 0.33.

The tuning profile of presynaptic cells was set according to the following generalized normal distribution: 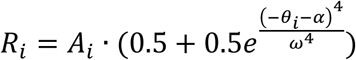, where *Ri* is the response from presynaptic cell *i*, α is the angle of stimulus direction, ω is the width of the presynaptic tuning profile (90°), and *Ai* is the response amplitude factor drawn at random for each presynaptic cell *i* before simulation run from a normal distribution with a mean of 4 and a SD of 4. Gaussian noise was implemented by changing the amplitude of each presynaptic cell individually by a value drawn at random for each trial from a normal distribution with the mean at 0 and variance equal to the noise-free presynaptic amplitude response. Directional classification errors were modeled by drawing a value from a normal distribution with a mean of 0 and SD of 45 for each presynaptic cell; this value was then added to the PD of each presynaptic cell and the tuning profile recalculated (Poleg-Polsky et al., 2018). This procedure effectively shifted the entire tuning profile of each presynaptic cell by a random value independently across trials.

### Noise correlations

Presynaptic population activity correlations in a trial were modeled iteratively over all 360 possible directions, in 20-degree advancements. In each iteration, a noise value was drawn as above for the corresponding direction angle (*ϑj*). The effect on the presynaptic responses in cell *i* was computed as a function of the distance of the cell’s PD from that angle: 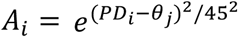. Next, a random value was drawn from a normal distribution whose variance was set by the noise level. Last, the amplitude/PD of the tuning profile of each cell was then changed by *Ai* multiplied by this random value for the ‘magnitude’/’directional’ noises respectively. This procedure guaranteed coordinated response fluctuations in cells with similar tuning profiles (Moreno-Bote et al., 2014; Pitkow et al., 2015; Kohn et al., 2016; Zylberberg et al., 2017).

To convert between presynaptic response amplitude and firing rate, the amplitude of each presynaptic cell was rounded to the nearest integer. This value indicated the number of presynaptic spikes from that cell. The temporal pattern was set by NEURON’s *negexp* function, with a mean interval between spikes of 20 ms. In addition to stimulus-mediated presynaptic activity, all presynaptic cells had a random firing rate of 4 Hz to emulate synaptic activation observed in the cortex. Where simulated, background synapses were activated with a random firing pattern; their average firing rate was 10 Hz.

### Binary discrimination

In these simulations, the presynaptic population was modeled as in the direction selectivity case. Only preferred and null (180° and 0° respectively) directions were probed. Classification errors were introduced as in the simple model (Figure 1) by randomly flipping signal identity in ∼15% of the presynaptic cells.

### Analysis

The number of stimulus evoked spikes for each presynaptic cell was measured within a 100 ms-long period that commenced at stimulus presentation (500 ms after simulation initialization). Fano Factor was computed as the ratio between the 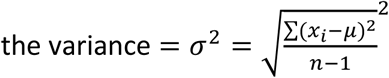 and the mean (µ) firing rates across cells within the 100 ms period. The skewness was computed as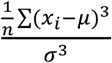, where *n* is the number of cells. All statistical analysis was performed in Igor7 (Wavemetrics).

The presence of dendritic spikes was measured as prolonged (>30 ms) depolarization above -30 mV in the activated dendritic location. Mutual information was measured as the entropy of signals across all stimulated directions minus the summed entropy of individual directions. Unless noted otherwise, error bars represent standard deviations.

The code used to generate this data can be accessed from https://data.mendeley.com/datasets/2rx82zw23n/draft?a=0cab0c63-a9cb-41f1-bae8-35814c2ebce3

## Results

### Noise sensitivity in linear vs. active dendrites

To illustrate the dependence of synaptic integration on compartment size, consider a binary discrimination task where we assume that a neuron is presented with two stimuli such that the preferred stimulus (to which the neuron should respond) is represented by 1 AU firing rate of the presynaptic population and a null (distractor) stimulus by 0 AU (**Figure 1 A**). We also assume that the trial-to-trial presynaptic input intensity follows a Gaussian (normal) distribution with a variance of one, giving a Fano Factor (variance/mean) of 1 for the preferred input, a realistic level of noise in the brain (Smith et al., 2013; Zhu et al., 2015; Zylberberg et al., 2016). We now ask how synaptic integration affects the ability of the postsynaptic cell to differentiate between the two stimuli in the presence of noise.

The precision of the encoding is diminished by the overlap in the distributions of presynaptic signals (**Figure 1 A**, right). The intersection between the signals is undesired because it reduces the ability of an ideal observer to discriminate between the stimuli. Consequently, the goal of a denoising algorithm is to reduce this overlap between population activity distributions – with the optimal outcome of non-overlapping stimuli representations. For such normal presynaptic population activity distribution, an acceptable denoising approach is a linear integration over the inputs. The trial-to-trial output of this algorithm also follows Gaussian distribution, which has the same mean as the original input but a narrower width [reduced from the original variance proportionally to the square root of the number of samples (**Figure 1 B, C**)]. Given a sufficient sample size, linear integration reliably distinguishes between the inputs; in the example shown in **Figure 1 B**, a linear integrator operating on 200 inputs produces a nearly perfect separation between preferred and null stimuli representations.

In contrast to the whole-cell integrator, Gaussian noise structure degrades the reliability of signal processing in active dendrites with a realistic integration compartment size of 10 inputs (**Figure 1 C**). The averaging capacity of such dendritic subunits is not enough to eliminate the overlap between dendritic signal representations – resulting in erroneous amplification of the null stimulus or a lack of dendritic spiking in response to the preferred drive (**Figure 1 C**). Unreliable synaptic integration in active dendrites in the presence of a high-variance input distribution reveals a potential computational weakness of dendritic subunits (**Figure 1 C**).

While dendritic processing is challenged by noise-corrupted signals in this example, nonlinear dendritic computations do not necessarily require low-noise conditions. Instead, active dendrites may be particularly sensitive to high-variance noises but perform well in the presence of other noise modalities. Recent empirical and theoretical studies have shown that stochastic Gaussian noise is not the only, and likely not even the most prevalent noise structure in the brain (Beck et al., 2012; Babadi and Sompolinsky, 2014; Pitkow et al., 2015; Gollnick et al., 2016; Kohn et al., 2016; Inagaki et al., 2019). Active dendrites may perform better when exposed to these alternative noise modalities.

### Dendritic spikes enhance signal estimation from asymmetric presynaptic distributions

While Gaussian distribution has been traditionally used to describe the trial-to-trial variability in the brain, most physiological responses to sensory stimulation actually follow asymmetric distributions (Buzsaki and Mizuseki, 2014). Population noise can take many forms, as I will show below. An important consequence of the asymmetric population variations is that the arithmetic mean no longer accurately describes noise-free input activity. This mathematical property reduces the denoising effectiveness of linear postsynaptic decoders because the (averaged) postsynaptic responses in the presence of noise diverge from the noise-free postsynaptic activity.

Simple examples of asymmetric response distributions to stimulation can be found in the primary auditory cortex that is stimulated with pure tones (DeWeese et al., 2003) or active touch encoding in the somatosensory cortex (Hires et al., 2015). Neurons in these cortices often respond with a single spike to their preferred stimulus and remain mostly silent otherwise (null trials; **Figure 1 D**). The behavior with respect to stimulation is almost binary and resembles processing in synthetic digital systems. A typical communication error in computers is a (mis)classification error, also known as a bit encoding error, which manifests as a mismatch between the transmitted and the received signals (Shannon, 1948). Similar classification errors are present in the brain – some neurons do not fire action potentials during the presentation of the preferred stimulus, and a non-zero rate of background activity implies that some cells will fire during the null trials (**Figure 1 D** left; (DeWeese et al., 2003)). The resulting distributions of firing responses are skewed in the opposite direction for the preferred and null responses (**Figure 1 D**, right). While most cells respond with a correct firing intensity to the stimulus in a given trial, the minority of cells that make classification errors pull the mean population activity towards the opposite response. The effect on the activity in the postsynaptic cell is significant, as the convergence of stimuli representations increases linearly with the percent of the misclassifying cells (**Figure 1 E**).

In contrast to the Gaussian noise, pooling across multiple presynaptic cells does not reduce the negative impact of classification errors (**Figure 1 E**) indicating that linear integration is not the optimal strategy to address misclassification. Instead, this noise structure necessitates a denoising mechanism that is based on a decision threshold – such as the threshold formed by regenerative dendritic spikes (**Figure 1 F**; (Hires et al., 2015)). These theoretical considerations suggest that the function of dendritic units is impacted by classification errors to a lesser degree than the computations in linear dendrites, perhaps enhancing neuronal ability to reliably process this noise modality.

### Distinct noise tolerance of linear vs. active dendrites revealed in a multi-compartmental model of a cortical cell

Studying the conceptually distinct sensitivities to Gaussian noise and classification errors of linear and active dendrites requires making observations of the input-output signal transformation in the postsynaptic cell. Although recent advances in high-resolution imaging techniques and electrophysiological recordings will likely be able to accurately describe the dynamics of the inputs to a cell at a single synapse resolution and the resulting postsynaptic output, such data are not yet available. For this reason, I took a modeling approach: I simulated signal processing in a detailed multi-compartmental NEURON model of a layer 2/3 cortical pyramidal cell in the visual cortex (see methods). This cell type was chosen because its basic biophysical properties and physiological function are both well understood (Larkum et al., 2007; Branco et al., 2010; Petersen and Crochet, 2013; Smith et al., 2013; Palmer et al., 2014; Ujfalussy et al., 2018).

To approximate a linear synaptic integration, I distributed α-amino-3-hydroxy-5-methyl-4-isoxazolepropionic acid receptor (AMPAR)-containing synapses randomly over the entire dendritic tree of the postsynaptic cell (**Figure 2 B**). To model synaptic integration in computational compartments formed by dendritic spikes, synaptic inputs containing both AMPAR and NMDARs were clustered on a single branch (**Figure 2 G**). NMDARs, which help to mediate glutamatergic transmission in many neurons, can trigger NMDA spikes in otherwise relatively passive dendrites (Polsky et al., 2004; Gordon et al., 2006; Major et al., 2008; Antic et al., 2010; Branco and Hausser, 2011; Chalifoux and Carter, 2011; Katona et al., 2011; Lavzin et al., 2012; Major et al., 2013; Palmer et al., 2014; Kumar et al., 2018)

**Figure 2:**
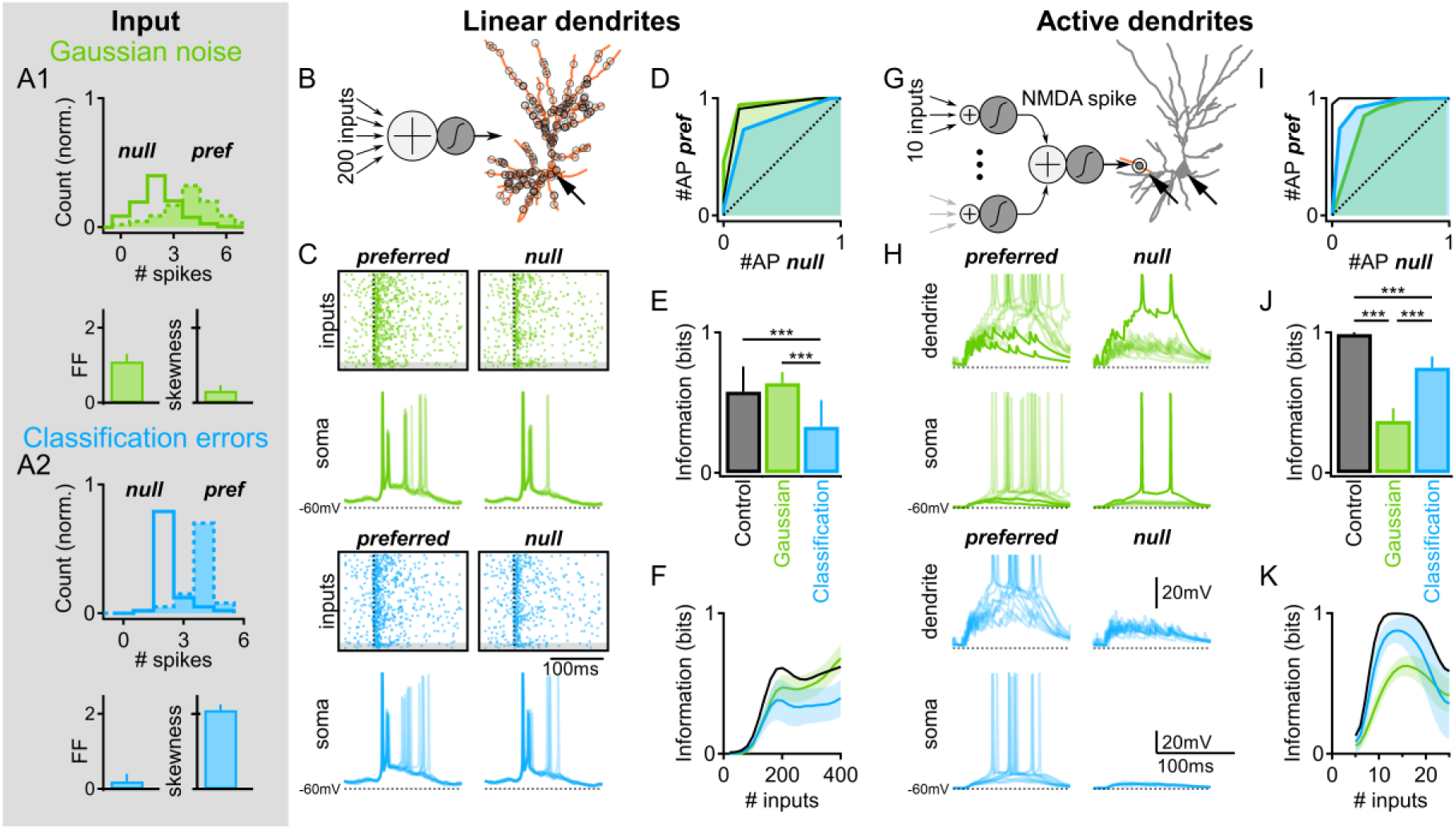
Distinct noise tolerance rules for linear vs. active dendrites in a reconstructed cortical layer 2/3 pyramidal cell. (A) Example presynaptic population dynamics with Gaussian noise (A1, Fano factor (FF) = 1) and classification errors (A2, error rate=15%). The mean population response to the preferred / null stimuli were 4 and 2 action potentials respectively. The baseline firing rates (4Hz) were similar across the simulated inputs. Top, the distribution of population firing rates. Bottom, Fano factor and the skewness calculated from the presynaptic distribution. (B-F) Synaptic integration in linear dendrites simulated by 200 randomly distributed AMPAR synapses (stimulated branches marked in red, B). (C) Top, example presynaptic spike trains in a single trial. Dotted line marks the time of stimulus onset. Bottom, postsynaptic voltages at the soma (marked with an arrow in B) in 10 representative trials in the presence of a Gaussian noise (green) or classification errors (blue). (D) Receiver Operating Characteristic (ROC) curve computed by plotting the normalized number of postsynaptic spikes for the preferred vs. the null stimuli presentations in 1000 simulation trials. Color coding as in (A); black, noise-free simulation. (E) The mutual information calculated from the postsynaptic output. ***p<0.001, n=1000 trials, ANOVA followed by Tukey test. (F) as in (E) as a function of the number of presynaptic inputs, shades - SD. (G-K) As for B-F for dendritic computational subunits engaged by activation of 10 clustered AMPAR/NMDAR synapses on the dendrite marked in red (G), which sampled a random subset of the inputs (C, gray). Arrows in G mark the dendritic and somatic recording locations.

Similar input statistics were presented in both integration schemes to ensure reliable comparison of noise tolerance rules in linear and active dendrites. The identity of the stimulus was encoded by the firing rates of the presynaptic population (**Figure 2**, methods). In the absence of noise, presynaptic cells spiked with twice as many action potentials in response to the preferred stimulus vs. the null stimulus (on average producing 4 and 2 spikes respectively). For a Gaussian population activity, the number of spikes for each presynaptic cell was drawn independently from a normal distribution with a variance = mean (**Figure 2 A1**). To model classification errors, 15% of the presynaptic cells were chosen at random to fire with the intensity of the opposite signal (**Figure 2 A2**). In addition to stimulus-evoked firing, all presynaptic cells had a baseline firing rate of 4 Hz.

To characterize postsynaptic noise sensitivity, I used the Receiver Operating Characteristic (ROC) analysis to compute the discrimination ability of a linear observer in the presence of noise. The normalized area under the ROC curve can vary between 0.5 - no discrimination and 1 - perfect performance. The area under the ROC curve measured from the somatic firing of the model with linear dendrites was not affected by the introduction of the Gaussian noise (from 0.9±0.05 in control to 0.92±0.06 with noise, **Figure 2 D**, green). As predicted by the conceptual model, this noise structure had a pronounced effect on synaptic integration in active dendrites; the area under the ROC curve diminished from 0.99±0.01 to 0.78±0.07 **Figure 2 I**, green). Similar results were observed when the information content of the postsynaptic output was measured with a mutual information metric. Mutual information is a useful indicator of the uncertainty between stimulus encoding and the output of a cell. The information is typically measured in bits, which specify the reduction in the number of yes/no questions needed to identify the stimulus after observing the outcome. The peak information in a binary discrimination task is 1 bit per trial, and a random input conveys zero bits. Gaussian noise did not change the information rate in linear dendrites (Figure 2 E, black vs. green), but dramatically reduced the performance of active dendrites (0.37±0.09 bits vs. 0.99±0.01 bits in control, **Figure 2 J**; p<0.001, n=1000, ANOVA followed with Tukey test).

Next, I measured the effect of classification errors on integration in both linear and active dendrites. The area under the ROC curve and the mutual information calculated from the postsynaptic firing were significantly degraded by classification errors in linear dendrites (**Figure 2 D, E**, blue) - despite signal transformation by somatic threshold nonlinearity (**Figure 2 C**). In contrast to the pernicious effect of classification errors on linear integration, this noise had a relatively mild effect on synaptic integration in active dendrites. Both the area under the ROC curve (0.92±0.05; **Figure 2 I**, blue) and the postsynaptic mutual information (0.75±0.08; **Figure 2 J**, blue) were significantly higher compared to the corresponding metrics in the presence of Gaussian noise (p<0.001, n=1000, ANOVA followed with Tukey test).

These results, observed for a wide range of stimulation intensities (**Figure 2 F, K**), and imposed noise levels (**Figure 3**) echo the enhanced tolerance of dendritic compartments to misclassification errors proposed by the conceptual model (**Figure 1**).

**Figure 3:**
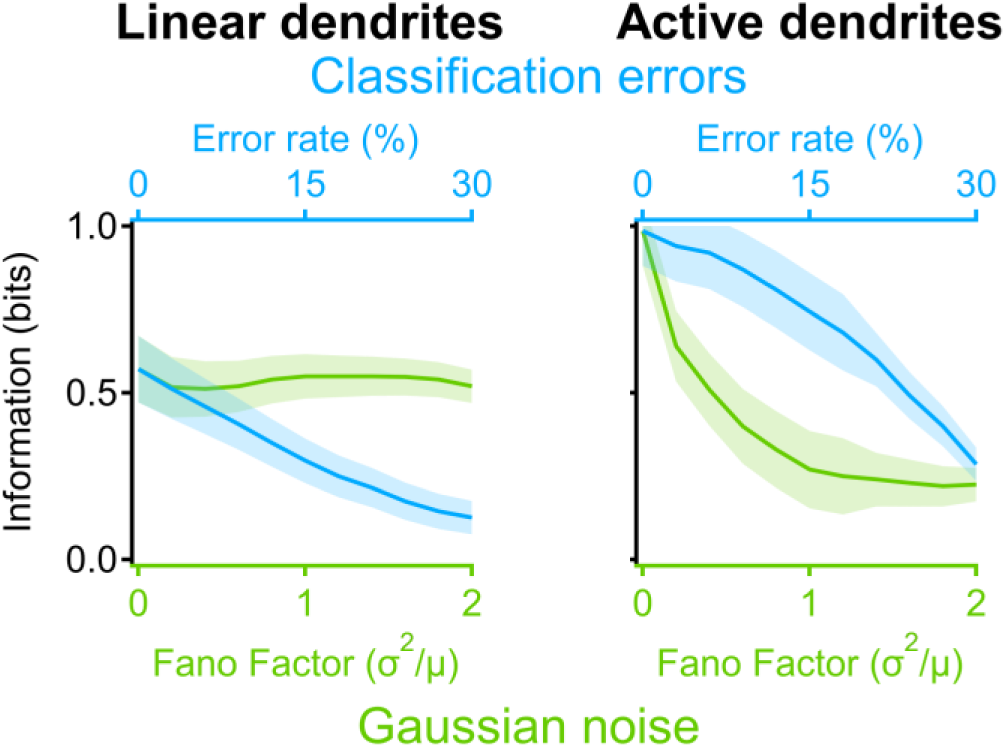
Performance in linear and active dendrites as a function of noise levels. Information rate per trial (mean ± SD), measured from the number of somatic spikes as a function of Gaussian noise intensity (green) or classification error rate (blue). Other model parameters were as in Figure 2.

### Dendritic computations in the presence of background synaptic activity

In the simulations presented so far, the postsynaptic cell was stimulated with signal-bearing synapses only. However, cortical cells in-vivo are exposed to a seemingly random background synaptic bombardment (London et al., 2010; Petersen, 2017). Variable membrane potentials mediated by the background drive can directly affect dendritic spike generation (Major et al., 2008; Lavzin et al., 2012; Smith et al., 2013). Can dendritic computations remain reliable in these challenging conditions?

To assess the sensitivity of dendritic processing to a more realistic synaptic environment, I added a signal-independent synaptic background. This background activity consisted of excitatory and inhibitory synapses distributed randomly over the dendritic tree (**Figure 4 A**, methods). Activation of these inputs depolarized the cell, increased the trial-to-trial variability of the resting membrane potential and promoted background postsynaptic firing (**Figure 4 B**) (Fellous et al., 2003). Yet, even at high levels of background activity, the postsynaptic cell was able detect, and signal the occurrence of the preferred stimulus (**Figure 4 B, C**). As before, computations in active dendrites were robust to classification errors, while linear dendrites were more informative during Gaussian noise (**Figure 4 C**).

**Figure 4:**
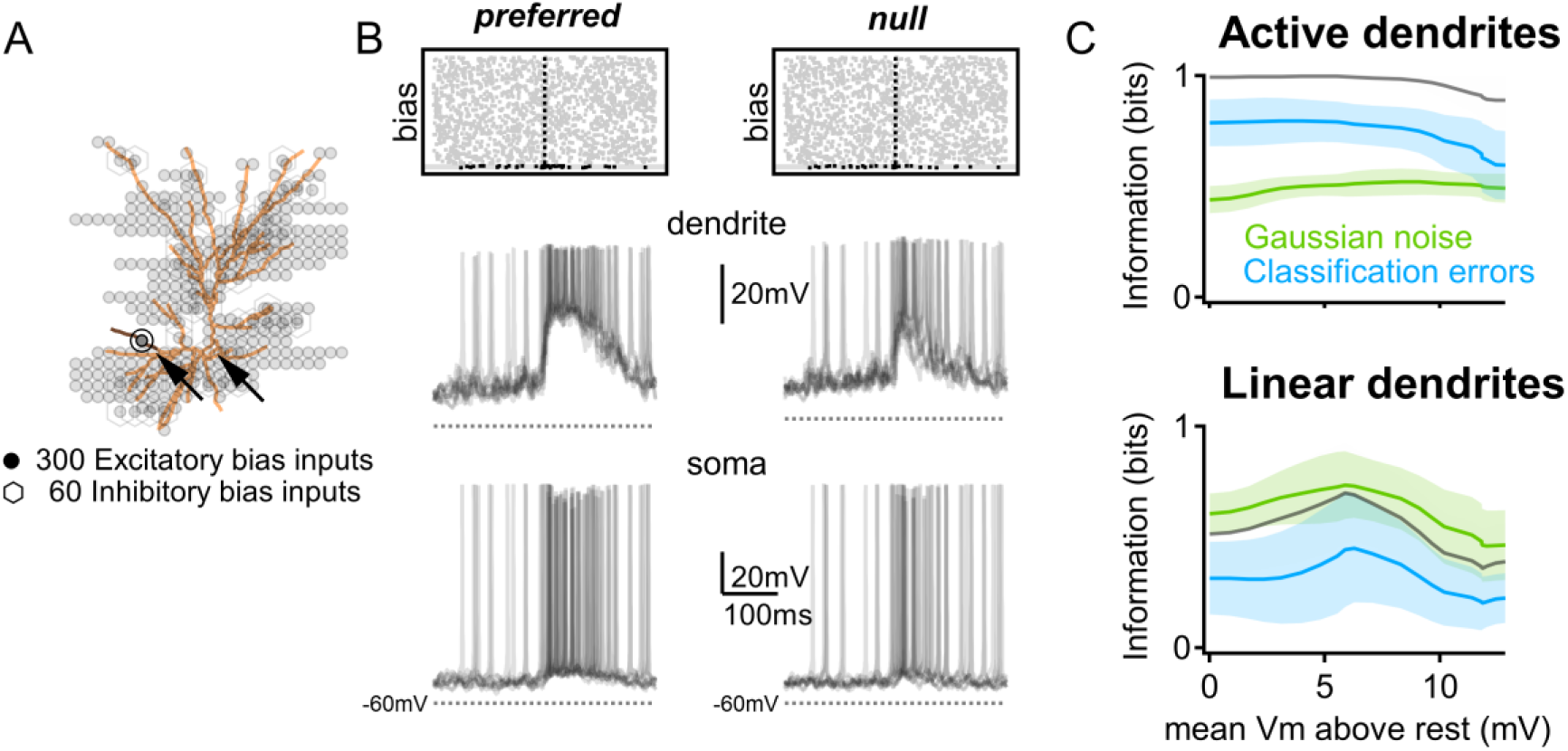
Active dendrites retain their performance in the presence of background activity. (A) Schematic position of the signal-mediating AMPAR/NMDAR synapses located on a single branch as in Figure 2, activated over a background of a randomly distributed 300 excitatory and 60 inhibitory inputs. (B) Top, example spiking activity in signal bearing (black) and background (‘bias’, gray) inputs. The dotted line indicates the time of stimulus presentation. Bottom, representative responses of the postsynaptic cell in 10 trials. Dendritic and somatic recording locations are marked by arrows (A). (C) The calculated mutual information as a function of the postsynaptic depolarization inflicted by background activity in models with active (top) or linear (bottom) dendrites. Black, noise-free simulation. Shades - SD. n=1000 repeats for each stimulus/condition. Note that the background activity does not significantly diminish the degree of information extracted by the postsynaptic cell over most bias levels.

### Distinct noise tolerance of linear vs. active dendrites in the computation of direction selectivity

Can the effects of noise on postsynaptic computations be generalized to more sophisticated computational tasks than a binary discrimination? To explore this question, I examined how noise affects direction selectivity (DS), a classic neuro-computation that is commonly used to analyze the impact of presynaptic signal variability on neuronal responses (Cafaro and Rieke, 2010; Franke et al., 2016; Kohn et al., 2016; Zylberberg et al., 2016). Dendritic computational subunits participate in the computation of direction selectivity (Oesch et al., 2005; Lavzin et al., 2012; Sivyer and Williams, 2013; Smith et al., 2013; Trenholm et al., 2014; Wilson et al., 2016), therefore, DS represents a relevant task to compare between synaptic integration in linear vs. active dendrites.

I began probing directional information processing in the presence of noise by assigning directional tuning to presynaptic neurons (**Figure 5 A**, black). The amplitude and the direction of the preferred response varied between presynaptic cells (methods). The Gaussian noise was modeled as before (**Figure 5 A1**, green). To account for multiple directions of stimulation, I implemented direction misclassification (directional errors) by shifting the directional preference of presynaptic cells (**Figures 5 A2**, blue): in each simulation trial, I rotated the entire directional tuning curve for each presynaptic cell by a random value taken from a normal distribution with a standard deviation of 45° (Poleg-Polsky et al., 2018). This manipulation mirrors labile directional tuning found experimentally in the cortex (Felsen et al., 2002; Benucci et al., 2013; Jeyabalaratnam et al., 2013) and other brain regions (Rivlin-Etzion et al., 2012).

**Figure 5:**
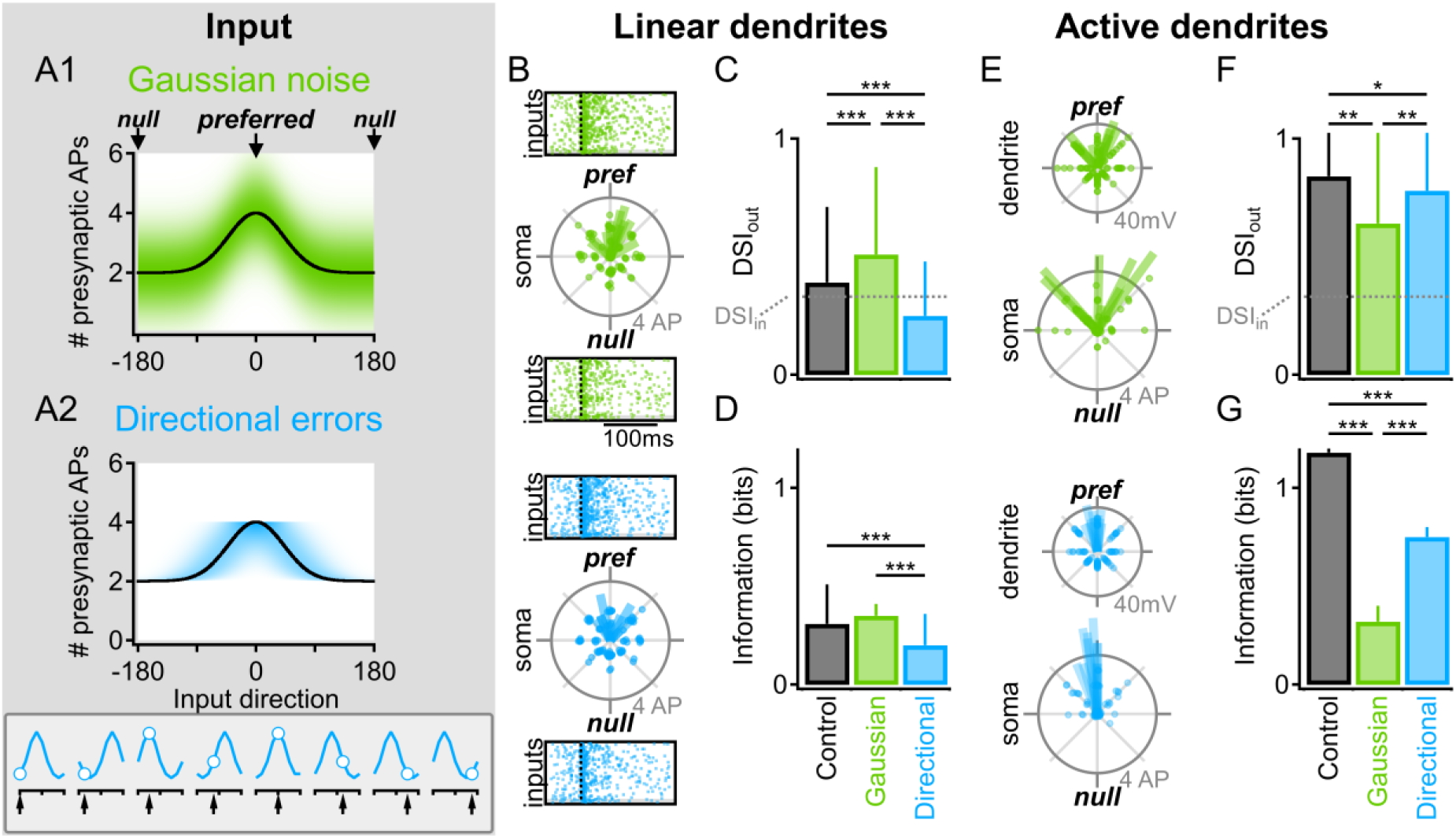
Distinct noise sensitivity rules in linear vs. active dendrites in the computation of direction selectivity. (A) Illustration of directional noise structures used in the model. (A1) Example tuning profile (firing rate vs. direction of the stimulus) of a single presynaptic cell in the presence of a Gaussian noise, implemented as fluctuations along the vertical axis. Shades of green indicate the response probability in the presence of noise (Black, noise-free condition). (A2) Directional tuning in the presence of classification errors. The noise was introduced as shifts in the directional profile along the bottom (direction) axis. Bottom, example construction of the presynaptic response from labile directional tuning (solid blue) in 8 individual trials. Arrows mark the probed direction. (B-D) Synaptic integration of directional information in linear dendrites. (B) Polar plots of the number of postsynaptic action potentials vs. the stimulated direction in the presence of population noises. Lines indicate the vector sum of responses in a stimulation round composed of 8 directions. Insets, representative presynaptic spike trains in a single trial. Dotted line marks the time of stimulus onset. (C-D) DSI (C) and mutual information (D) transmitted by the somatic firing of the postsynaptic cell in a stimulation round. (E-G) As in (B-D) for active dendrites. Dendritic directional tuning in (E) was computed from the peak potentials in the active dendrite. *p<0.05, **p<0.01,***p<0.001, n=1000 rounds, ANOVA followed by Tukey test. Error bars represent SD.

Next, I recorded dendritic potentials and somatic spikes in the presence of Gaussian noise and directional errors (**Figure 4 B-G**) and analyzed the sharpness of the postsynaptic directional tuning with Direction Selectivity Index 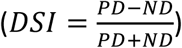 and with mutual information, calculated from the somatic firing for 8 directions of stimulation. The DSI output in the linear dendrites model was only marginally better in the noise-free case than the DSI of the input (0.39±0.32 vs. 0.33, **Figure 5 C**). Direction selectivity in the presence of Gaussian noise was higher than in control (DSI=0.51±0.37), but reduced to 0.25±0.23 when directional errors were introduced (**Figure 5 C**; p<0.001 between the three simulated conditions, n=1000, ANOVA followed by Tukey test). Similar trends were observed in the information content of the postsynaptic firing response (**Figure 5 D**).

An important impact of active dendritic processing was on the DSI levels and the information content of the postsynaptic cell, both of which increase well above the linear model (DSI=0.84±0.22 and 1.2±0.02 bits/trial respectively, **Figure 5 F, G**, noise-free condition, black). Similar to the binary discrimination task, the reliability of the computation was significantly impacted in the presence of Gaussian noise (**Figure 5 E-G**, green), but less so by directional errors (**Figure 5 E-G**, blue). These results indicate that the different noise tolerance rules for linear vs. active dendrites are apparent in salient computations involving graded signals.

### The impact of input correlations on noise tolerance of active dendrites

Despite the importance of the effect of independent (uncorrelated) fluctuations in the presynaptic population, the discussion of the effect of noise on active dendrites is incomplete without considering coordinated presynaptic activity. Therefore, here I focused on correlations in the shared response variability to repeated presentation of the same stimulus, often referred to as noise correlations, which are different from signal correlations that indicate preference for similar stimuli (Cohen and Kohn, 2011; Kohn et al., 2016).

The impact of correlated population code on information detection in linear dendrites is well understood (Moreno-Bote et al., 2014; Pitkow et al., 2015; Franke et al., 2016; Kohn et al., 2016; Zylberberg et al., 2016; Zavitz et al., 2017). The black line in **Figure 6 A** illustrates an example trajectory of a pair of direction-selective neurons, for diverse stimulus parameters in the absence of noise. This trajectory can be expanded into a multi-dimensional signal space for the entire population (**Figure 6 B**). A noise that pushes the population response away from this signal space is considered to be relatively benign, while noises that spread the responses along the signal space are thought to limit the precision with which the signal can be decoded (**Figure 6 A, B**). Thus, noise structures that mimic the signal being transmitted should have a disproportionally higher negative impact on the fidelity of information encoding by the postsynaptic cell.

**Figure 6:**
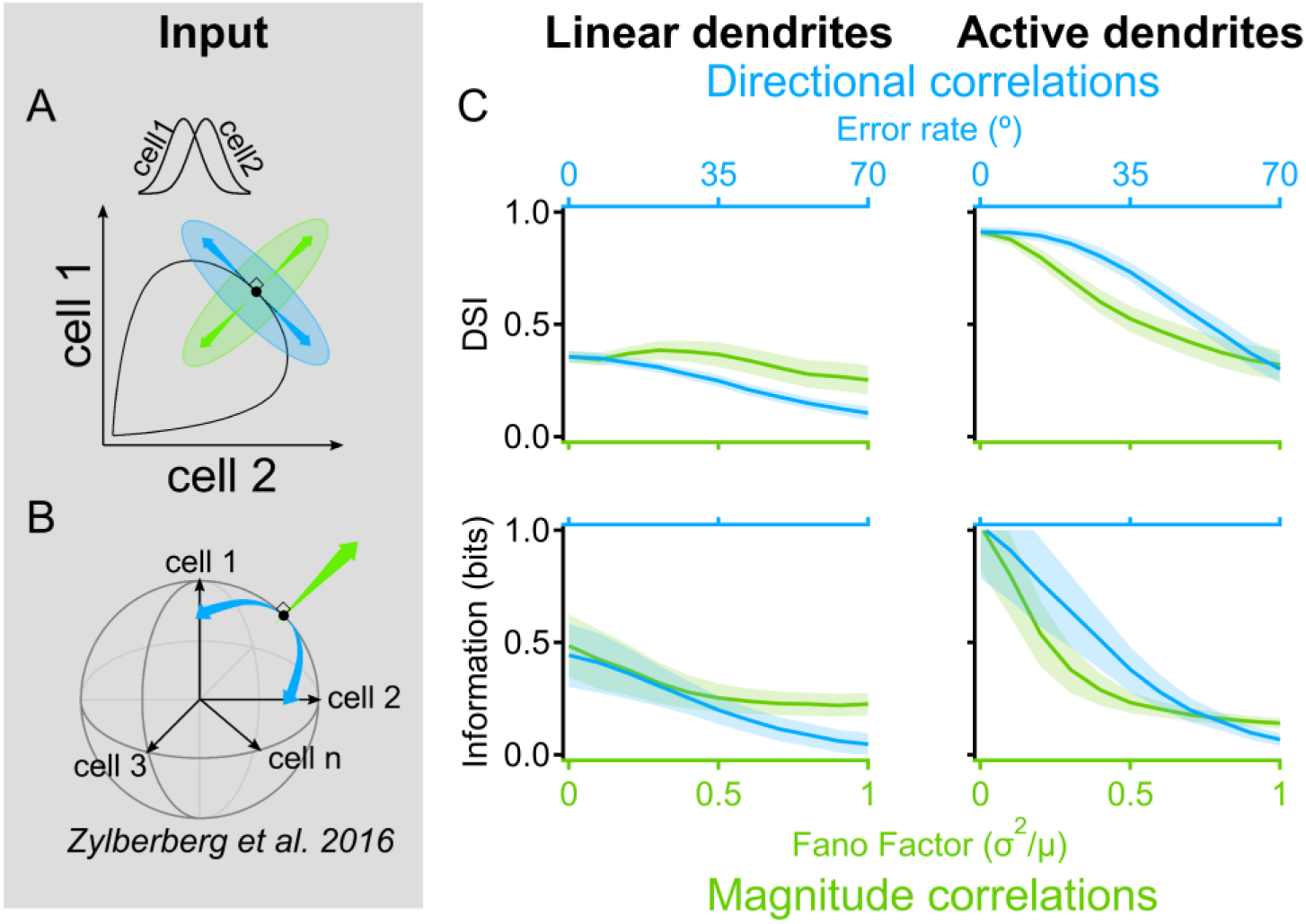
Signal-space noise correlations enhance information capacity of active dendrites. Population noise can be described by a vector in the n-dimensional space where each dimension corresponds to the activity of a single presynaptic cell. The trial-to-trial correlation structure between neuronal responses determines the amount of information that can be represented by the population. (A) Illustrated neural response space with two neurons with different directional tuning curves (inset). Black, the noise-free activity plotted for all possible stimulus angles. Green, a ‘magnitude’ noise structure that shifts the trial-to-trial response variability along a vector that points in the orthogonal direction from the population curve. Magnitude noise was shown to be well-tolerated in linear dendrites. Blue, ‘directional’ noise correlations, which translate the responses along a tangent to the curve, are known to diminish the information that can be extracted with linear integration (Moreno-Bote et al., 2014; Zylberberg et al., 2016). (B) Similar to (A) for *n* presynaptic cells. Directional correlations tend to vary the population activity along the signal space that mostly lies within the shell of a hypersphere (Zylberberg et al., 2016). (C) Postsynaptic DSI (top) and mutual information (bottom) vs. noise intensity in linear (left) and active (right) dendrites. Presynaptic tuning curves and correlations statistics remained similar across trials. Unlike linear dendrites, processing with dendritic spikes produced better DSI values and carried more information about the stimulus in the presence of directional correlations. Shades – SEM.

Given the difference in noise tolerance between linear vs. active dendrites, is it possible that noise correlations have different effects on the decoding abilities of dendritic subunits? To address this question, I modeled correlated activity in the presynaptic population. The geometric representation of the signals in the multi-dimensional space helped to design correlated noise structures. The signal space, or the activity of the presynaptic population, is largely restricted to the shell of a hypersphere, where each axis represents the mean activation of a single presynaptic cell (**Figure 6 B**; (Zylberberg et al., 2016)). ‘Directional’ correlations arise from similar directional errors in a sub-population of the inputs and follow the curvature of the signal space (**Figure 6 A, B**, blue). Fluctuations that are orthogonal to the signal space point in the radial direction (**Figure 5 A, B**, green (Zylberberg et al., 2016)). I simulated these ‘magnitude’ correlations by picking similar response values from a Gaussian distribution across neurons with similar directional preference (Moreno-Bote et al., 2014; Zylberberg et al., 2016). This algorithm provides an intuitive explanation for how the formulation of uncorrelated directional and Gaussian noises can be expanded to include correlations. Based on the differential effects of uncorrelated noise structures, it is likely that the function of active dendrites will be better maintained in the presence of directional correlations. To examine this prediction, I varied the intensity of the noises by changing the mean size of the noise vector - either as a function of the direction of the stimulus (directional noise) or the amplitude of the Gaussian noise distribution (magnitude noise; **Figure 6 C**). In line with the findings of previous studies (Pitkow et al., 2015; Franke et al., 2016; Zylberberg et al., 2016), the DSI and the information calculated from the firing output of the linearly integrating postsynaptic cell were both consistently higher for magnitude noise correlations (**Figure 6 C**, left). Importantly, this relationship was reversed for active dendrites, where shifts in population activity along the signal space were integrated better than the orthogonal, magnitude correlations across much of the tested noise range (**Figure 6 C**, right). This result supports the conclusion that differences in synaptic integration between linear vs. active dendrites found in independent population activity apply to correlated noise structures.

## Discussion

In this study, I examine how synaptic integration in active dendrites affects the robustness of neuronal computations in noisy environments. By analyzing simulated postsynaptic responses to different empirically characterized population activity patterns, I describe for the first time that neuronal computations with dendritic processing units do not follow the noise tolerance rules established in conventional population models. Simple conceptual models and biophysically realistic simulations demonstrate differences between linear dendrites vs. active dendrites, both in terms of denoising strategies, as well as robustness for discrimination of dissimilar noise structures (**Figure 7**).

**Figure 7:**
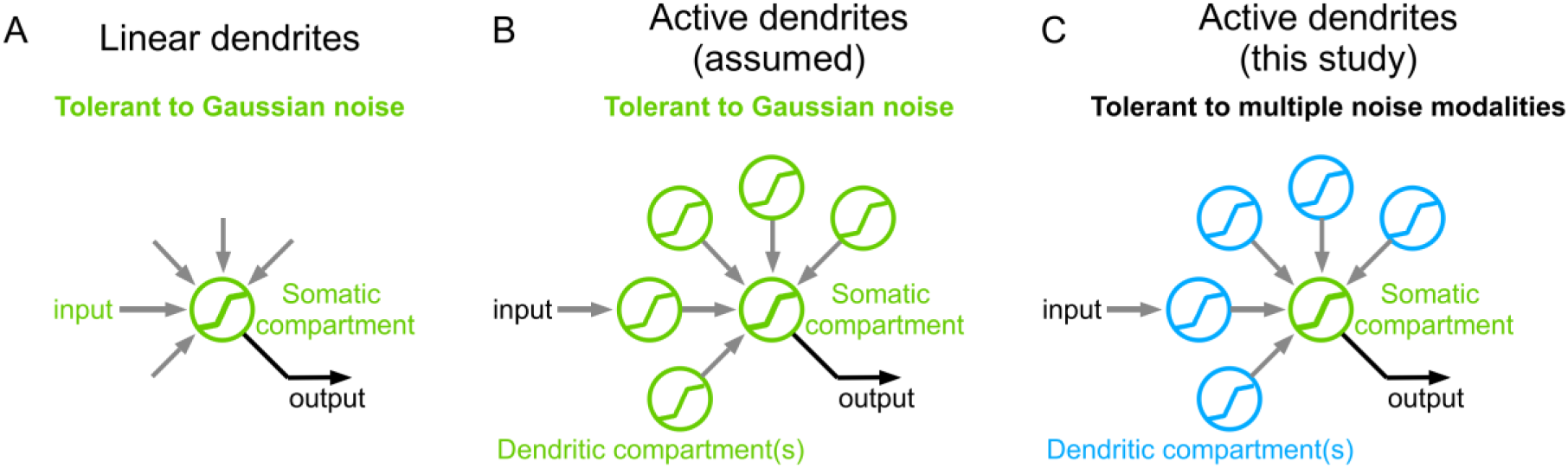
Summary of the main findings. (A) Traditionally, synaptic integration is considered to be linear and robust to stochastic changes in presynaptic input amplitudes (inset, green). (B, C) Real neurons often integrate their inputs nonlinearly in dendritic computational subunits. These subunits were assumed to have a similar noise profile to linear integrators (B). Here I report a distinct noise tolerance of active dendrites. While susceptible to high-variance noise, dendritic compartments excel in the presence of binary classification errors, labile directional tuning and noise correlations that limit the information that can be extracted with linear dendrites (C). Dendritic spikes can thus help expand the repertoire of computationally benign noise structures.

### Dendritic spikes are important for cortical processing

In mammals, various neuronal types possess the necessary dendritic machinery for focal dendritic spike generation (Manita et al., 2017). Dendritic spikes are potent mediators of somatic firing (Oesch et al., 2005; Grienberger et al., 2014; Palmer et al., 2014; Schmidt-Hieber et al., 2017). They contribute to neuronal computations in-vivo and correlate with perception (Lavzin et al., 2012; Smith et al., 2013; Takahashi et al., 2016; Wilson et al., 2016; Manita et al., 2017; Schmidt-Hieber et al., 2017). The calcium influx associated with a dendritic spike is an important mechanism for induction of synaptic plasticity (Gambino et al., 2014), linking dendritic regenerativity with learning (Kaifosh and Losonczy, 2016; Schiess et al., 2016) and memory formation (Sheffield and Dombeck, 2015; Guerguiev et al., 2017).

I have shown that computations in active dendrites consistently outperform linear signal integration (**Figures 2-6**). This finding resonates with the progressive enhancement of metrics for direction and orientation selectivity in the cortex that occurs in the presence of dendritic spikes. In mice, thalamic input to V1 is weakly tuned (Sun et al., 2016) similar to the presynaptic drive used in current simulations (**Figure 5**). The increase in cortical visual selectivity between processing layers is most pronounced in neurons that exhibit dendritic nonlinear input-output transformation (Lavzin et al., 2012; Smith et al., 2013; Ujfalussy et al., 2015; Wilson et al., 2016), [but see (Jia et al., 2010)]. Suppression of dendritic spikes in the cortex by hyperpolarization or intracellular perfusion of NMDAR blockers results in decreased subthreshold and suprathreshold selectivity, confirming that dendritic amplification mechanisms are crucial for synaptic integration (Lavzin et al., 2012; Smith et al., 2013). These multiple lines of investigation strongly suggest that reliable dendritic spike generation is essential for brain function.

Currently, the consensus is that by combining neural responses, the brain can average out the impact of noise and increase the signal to noise ratio (SNR). Linear input integration is especially useful in limiting stochastic fluctuations in synaptic signals – implying that a Gaussian noise source is relatively benign for signal decoding, but only when the sampled presynaptic population is large. As I have shown here, due to their relatively small integrative compartment sizes, dendritic spikes could be particularly sensitive to variability at the SNR levels that are common in the brain (Magee, 2000; Smith et al., 2013; Franke et al., 2016; Zylberberg et al., 2016; Zavitz et al., 2017).

### Evidence for non – Gaussian noise in the brain

While we intuitively assume that noise fluctuations follow a Gaussian distribution, noise representation in the brain is a fundamental open question in neuroscience. I have taken the question of ‘what is noise?’ one step further. A broad class of neuronal computations can be formulated as discrimination problems, where the network is tasked with sorting the input into distinct categories. Beck et al. (2012) provided compelling theoretical evidence to suggest that misclassification noise is practically unavoidable in complex nervous systems due to suboptimal inference (Beck et al., 2012). Misclassifications plague artificial neural networks; our conscious experience suggests that classification errors are common in biological brains. Instead of treating classification errors as computational failures, I propose to consider them as a noise source that decreases the accuracy of the computation.

Experimental results provide examples of classification errors. Cortical responses to sensory stimulation can be binary, composed of a single or no action potentials, as originally described by DeWeese et al. in the primary auditory cortex of the rat (DeWeese et al., 2003) and more recently seen in the mouse barrel cortex (Hires et al., 2015; Ranjbar-Slamloo and Arabzadeh, 2017). Neurons in these areas fire for most, but not all, repeated stimulation trials and often have a non-zero background firing rate in the absence of stimulation (DeWeese et al., 2003; Tao et al., 2016). Classification errors are harder to distinguish from the normally distributed fluctuations for stronger firing responses (DeWeese et al., 2003; Tao et al., 2016) and more complex stimulation paradigms (Poleg-Polsky et al., 2018). A useful criterion that can discriminate between different noise structures is symmetry. By definition, a Gaussian process elicits responses that are normally (symmetrically) distributed around the noise-free mean [**Figures 1-2**, (Poleg-Polsky et al., 2018)]. Thus, trial-to-trial response variability that is not symmetric is likely to arise from a different source. An example of a complex, non-stochastic signal variability in the brain was demonstrated by Gollnick et al. (Gollnick et al., 2016). The authors revealed a probabilistic cortical population encoding of whisker deflection intensity where the sensory input affected the likelihood of a neuronal response, but not the mean response intensity. Furthermore, the magnitude of whisker deflection could not be estimated correctly with a linearly operating ideal observer, in line with my findings that linear decoders fail with a non-symmetric input profile.

Despite the high trial-to-trial SNR levels in the presence of classification errors, linear mechanisms cannot average away this noise, irrespective of the size of the presynaptic population (**Figures 1-2**). The situation is different with dendritic computations because nonlinear synaptic integration utilizes spike threshold to increase the reliability of presynaptic input classification. In **Figures 1-4**, I showed how this process works for a binary decision process; the probability of dendritic spike initiation encodes the information in the case of a continuous variable (**Figure 5**). These results are consistent with, and expand on, previous theoretical observations that dendritic computations are reliable with high-SNR manipulations (Rhodes, 2008; Wu and Mel, 2009; Hawkins and Ahmad, 2016).

### Synaptic integration of correlated noise structures

Studies of neuronal population dynamics have primarily focused on network computations emerging from interactions between highly simplified, linearly integrating neurons (Seung and Sompolinsky, 1993; Zohary et al., 1994; Vogels et al., 2011), but see (Zylberberg et al., 2017). These investigations revealed that coordinated neuronal activity has a substantial impact on information encoding. Arguably, one of the most interesting findings is that correlations that change the variability of neuronal responses along the direction parallel to the multi-dimensional signal space distort the population activity in a way that cannot be averaged out, limiting the accuracy of stimulus decoding. This correlation structure is known as differential (Moreno-Bote et al., 2014), choice (Pitkow et al., 2015) or directional noise (Zylberberg et al., 2016). To understand why these correlations are information limiting, consider estimation of stimulus direction from responses perturbed by this noise structure. The activity of the population shifts from the noise-free position on the signal space towards another point on (or close to) that space, which by definition represents a noise-free response to a different stimulus direction. The estimation of an ideal linear decoder will deviate from the true stimulus, irrespective of the number of presynaptic inputs (Kohn et al., 2016). Thus, similar to uncorrelated noise structures, unreliable encoding of stimulus identity degrades the performance of a linear integrator. Because differential/directional correlations are prevalent in the brain, neural coding has been suggested to be inherently suboptimal (Pitkow et al., 2015).

My results support the opposite conclusion: directional correlations are a relatively benign noise structure for signal processing in small dendritic integration compartments. The quality of information transfer by a small number of correlated inputs is already mostly independent of the structure of the noise – at least for a linear decoder (Pitkow et al., 2015). As in the uncorrelated directional noise condition, the dendritic spike threshold provides a decision cutoff that allows the postsynaptic cell to categorize the stimulus. As a result, active dendrites do not directly help in addressing the question of what was the direction of the stimulus. Rather, they signal the probability that the stimulus was in the preferred, null or intermediate directions, which is likely to be more behaviorally-relevant information.

To conclude, my analysis of a binary discrimination task and both correlated and uncorrelated directional noises drew strong conclusions about information processing in active dendrites. Computations performed with dendritic spikes are necessary to enhance salient stimuli features, but the conventional wisdom is that synaptic noise should have a significant negative impact on the reliability of dendritic calculations. In contrast with this view, I present theoretical arguments in favor of the idea that dendritic computational compartments have robust tolerance to noise. I argue that active dendrites provide neurons with an optimal strategy to process noise structures that differ from the ‘benign’ noise types established for linear neurons. These conclusions characterize a key component of cortical signal processing and highlight the importance of studying the impact of noise in the context of postsynaptic integration, because signal transformation in dendrites can fundamentally change our understanding of how brains process sensory information.

## Acknowledgments

I thank John Gaynes, Gautam Awatramani, Jeffrey Diamond and Joel Zylberberg for helpful comments on the manuscript.

